# Predicting pain location from resting-state brain fMRI

**DOI:** 10.64898/2026.06.14.732139

**Authors:** Jennifer A. Cummings, Sharmila Majumdar, Andrew Bishara, Julian Motzkin, Ashish Raj, Prasad Shirvalkar, Jeffrey Lotz

**Author notes:** Contributing authors.

## Abstract

Low back pain is a prevalent issue with few reliable treatments. Although there is great variation in clinical presentation within the low back pain population, little is known about the neurobiological mechanisms underlying these differences. In this study, we sought to stratify chronic low back pain patients (N = 275) into phenotypes characterized by correlated patterns of resting-state brain activity and sensory abnormalities (pain, numbness, and pins and needles) indicated on hand-drawn body maps. Our cross-decomposition analysis yielded phenotypes that resemble previously documented mechanistic pain types, revealing distinct brain connectivity patterns associated with different clinical presentations. Our model was then used to predict pain body maps from fMRI data in a small novel dataset of chronic pain subjects, suggesting that these relationships may generalize to other chronic pain conditions. Our results support the utility of resting-state fMRI in understanding the heterogeneity of chronic pain, which may be leveraged to develop more targeted pain treatments.

## 1 Introduction

Low back pain (LBP) is the leading cause of disability worldwide, with approximately 90% of LBP patients classified as “non-specific” Wu et al (2020); Maher et al (2017). This diagnostic ambiguity, coupled with the limited utility of spine imaging findings, has led to numerous ineffective, invasive, and costly interventions Brinjikji et al (2015). Despite the biopsychosocial model’s growing adoption and the implementation of multidisciplinary treatment approaches, clinical outcomes remain modest Kamper et al (2014), underscoring the need to identify distinct pain phenotypes that could benefit from targeted interventions.

Neuroscience may provide some guidance toward this goal. A growing body of evidence associates chronic LBP (cLBP) with structural and functional changes in the nervous system when compared to pain free individuals Kregel et al (2015); Konno and Sekiguchi (2018); Li et al (2024). Changes have been shown to coincide with the transition from acute to chronic pain and may even predict someone’s likelihood to transition after injury Hashmi et al (2013); Baliki et al (2012). Additionally, studies have suggested that some neurobiological signatures may normalize after successful pain treatment Seminowicz et al (2011).

While numerous studies have investigated neurobiological differences between cLBP patients and healthy controls, significantly less work has focused on parsing sources of phenotypic variation within the cLBP population itself. Despite widespread adoption of the mechanistic categorization framework proposed by the International Association for the Study of Pain (IASP), the neurobiological signatures underlying these distinct clinical presentations remain poorly understood. The complexity of these signatures is evident across pain types: the thalamus shows inconsistent involvement in neuropathic pain, with studies reporting both hyper- and hypoactivity Garcia-Larrea and Peyron (2013); Alshelh et al (2016); both neuropathic and nociplastic profiles demonstrate higher neuroimmune activity in S1 and stronger S1-thalamus functional connectivity compared to nociceptive types Alshelh et al (2022); and nociplastic pain (recently adopted as a third IASP category) appears characterized by inhibited descending pathways and broad sensory processing abnormalities Harte et al (2018), making it particularly important for treatment planning Clauw (2024). These heterogeneous findings suggest that multivariate analytical approaches may be better suited to capturing the complex neurobiological patterns that differentiate pain mechanisms, ultimately aiding in more precise treatments for chronic pain patients.

In recent years, canonical correlation analysis (CCA) has risen to prominence for its ability to reveal complex relationships in multi-modal, high-dimensional data. CCA is a multivariate statistical method that maximizes the correlation between two or more datasets, making it uniquely suited for neuroscience datasets, which often contain many different types of data (e.g. neuroimaging, behavioral, genetic, etc.) from the same subjects Hotelling (1992); Wang et al (2020); Zhuang et al (2020); Mihalik et al (2022). Inspired by recent studies using CCA to stratify heterogeneity within a disease population, we hypothesized that the method may reveal hidden phenotypes in our cLBP cohort Drysdale et al (2017); Buch et al (2023).

While numerical pain scales like the visual analog scale (VAS) or numeric rating scale (NRS) would provide the most direct analogy to behavioral measurements used in previous CCA studies, these are insufficient for capturing the full dimensionality of the chronic pain experience van Boekel et al (2017). In contrast, self-reported pain maps contain a wealth of information but have long evaded use in research due to a lack of standardization and quantification practices. However, growing interest in the link between pain widespreadedness and central sensitization has brought a new focus on body maps as a measurement tool Brummett et al (2016); Scherrer et al (2021); Fitzcharles et al (2021). Body map data has already proven useful in guiding treatment planning, with those reporting a greater spatial extent of pain showing an increased likelihood to respond well to centrally acting therapies such as analgesics Clauw (2024). Thus, body maps seemed an ideal source from which we could mine measurements that would best capture the individual variability in our cLBP subjects.

The goal of our study was to identify neurological signatures corresponding to unique somatosensory profiles in a dataset of nearly 300 subjects with chronic low back pain. To that end, we developed a computational pipeline for extracting meaningful measurements from hand-drawn body maps of three sensory abnormalities (pain, numbness, and pins and needles) while maintaining a balance between sensitivity and parsimony. We then applied regularized CCA to identify axes of common variability between resting-state functional connectivity data and these body map-derived metrics, termed “brain-body dimensions”, which were validated in held-out test sets using a bootstrapped cross-validation procedure. Some of these dimensions show remarkable correspondence with the established mechanistic pain categories, providing new insight into the neurobiological signatures of these phenotypes. The model was also shown to predict pain reports in a novel dataset of chronic pain patients, suggesting that they may not be unique to LBP. Finally, we investigated the relationship between these brain-body dimensions and several clinical, demographic, and psychosocial risk and prognostic factors.

To our knowledge, this study represents the first investigation to show an association between whole-body somatosensory patterns and brain connectivity recorded at rest. Our results provide new evidence supporting a link between chronic pain and brain network reorganization measurable in absence of any task or pain stimulus. These findings also underline the complexity of cLBP, showing that no single brain connectivity feature or somatosensory profile can describe all LBP subjects. We hope that these results will encourage further efforts toward stratifying the heterogeneous LBP population into clinically meaningful phenotypes with the ultimate goal of providing personalized effective treatments.

## 2 Results

In summary, we performed regularized CCA (rCCA) to identify dimensions of common variation between resting-state brain connectivity and whole body somatosensory patterns identified from patient-reported body maps. Parameter optimization was performed using k-fold cross-validation, yielding a final solution of 7 canonical components, which we refer to as “brain-body dimensions”. All 7 dimensions were found significant in held-out validation sets using bootstrapped permutation testing (Supplementary Figure B1). Model weights were then used to predict pain maps for a set of previously unseen subjects, suggesting an ability to identify the spatial distribution of chronic pain symptoms from brain activity recorded at rest.

### 2.1 A subset of brain networks predicts somatosensory symptoms

A subset of 350 resting-state functional connectivity features were selected as input to the rCCA model, all of which are represented in Figure 1. These were selected for their strong correlations with at least one group-level body map pattern (details in 4.6); thus, each plot provides a different view into the neural correlates of pain and sensory abnormalities in this cohort. The chord plot (top left) shows that features largely involved connectivity between a subset of cortical and subcortical networks. Specifically, connections between the somatomotor and frontoparietal networks are common, as well as connections between subcortical regions and parts of the default mode network (DMN) and dorsal attention networks. The glass brain (top right) shows specific nodes corresponding to regions of the Brainnetome atlas Fan et al (2016). Node sizes correlate with degree, meaning that larger nodes are involved in more of the edges chosen by the model. Three nodes are especially highly represented, each involved in more than 5% of the 350 features. These are the left and right homologues of region A2 of the postcentral gyrus, corresponding to the primary somatosensory cortex (S1), as well as the left region A20cl of the inferior temporal lobe. Finally, the bar plot (bottom) shows the summed degree of all nodes within each anatomical region, colored by node network. Here we see that a majority of connections involve regions of the parietal lobe and the thalamus.

**Fig. 1.**
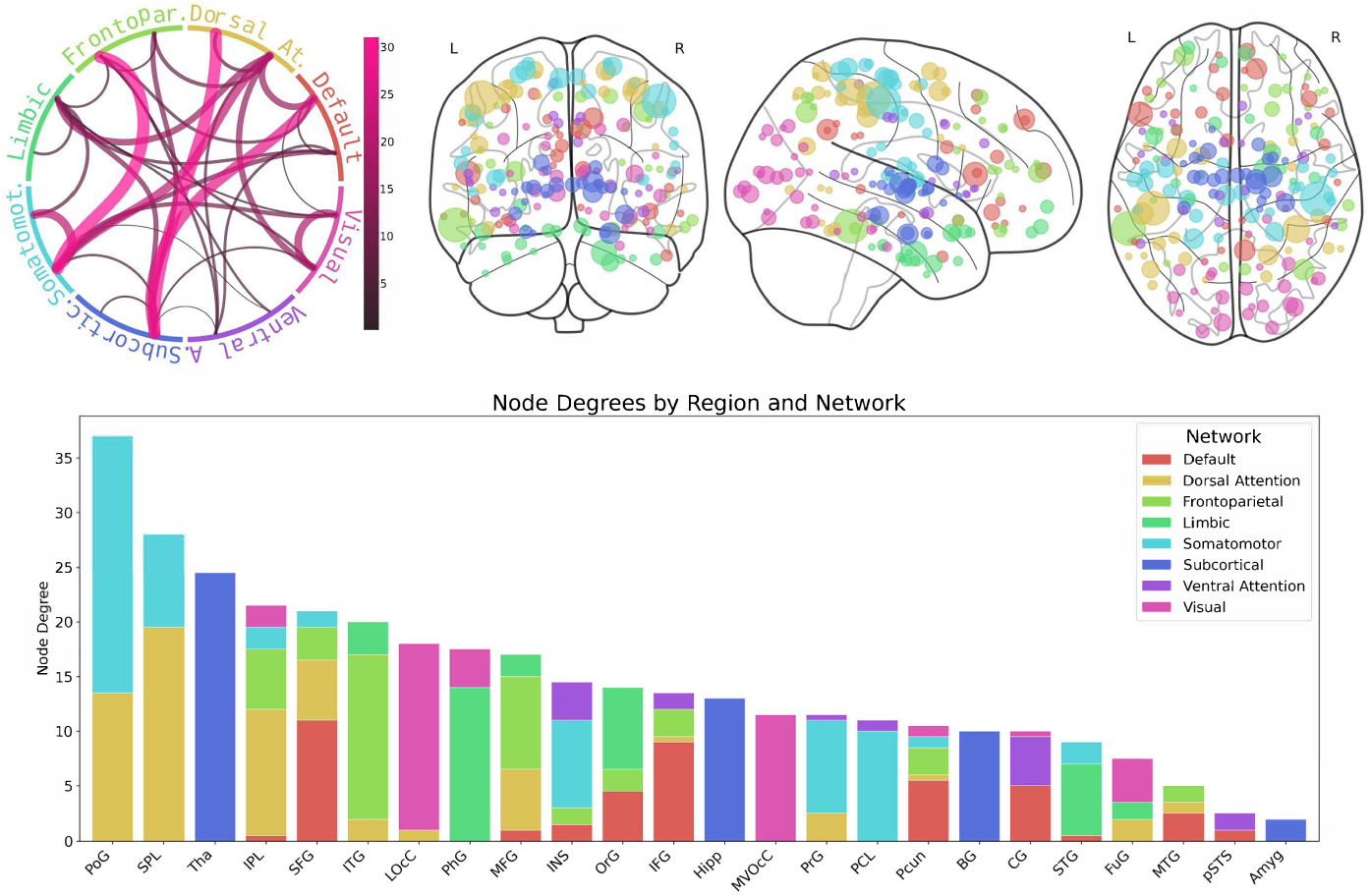
All 350 resting-state FC features used in final model. Data are represented in three ways to aid in interpretation. Chord plot (upper left) shows network membership of all edges with line color and thickness indicating number of occurrences. Glass brain (upper right) shows anatomical locations of nodes, with nodes colored by network membership and sized by degree. Bar plot (bottom row) shows summed node degree by network and region in descending order. SFG: Superior Frontal Gyrus, MFG: Middle Frontal Gyrus, IFG: Inferior Frontal Gyrus, OrG: Orbital Gyrus, PrG: Precentral Gyrus, PCL: Paracentral Lobule, STG: Superior Temporal Gyrus, MTG: Middle Temporal Gyrus, ITG: Inferior Temporal Gyrus, FuG: Fusiform Gyrus, PhG: Parahippocampal Gyrus, pSTS: posterior Superior Temporal Sulcus, SPL: Superior Parietal Lobule, IPL: Inferior Parietal Lobule, Pcun: Precuneus, PoG: Postcentral Gyrus, INS: Insular Gyrus, CG: Cingulate Gyrus, MVOcC: MedioVentral Occipital Cortex, LOcC: Lateral Occipital Cortex, Amyg: Amygdala, Hipp: Hippocampus, BG: Basal Ganglia, Tha: Thalamus.

### 2.2 Brain-body dimensions resemble mechanistic pain categories

Each of the 7 components identified through rCCA represents a “brain-body dimension” in which somatosensory patterns are linearly correlated with resting-state connectivity patterns. All components are strongly correlated (*p* range 0.77-0.83) and significant in held-out test sets compared to shuffled data (see Appendix B). A figure showing the brain and body features associated with each component is shown in the Extended Data (Appendix C; Figures C2 - C6). Here, we focus our analysis on the first two components based on their distinctive alignment with previous literature. Specifically, we found that the body map patterns associated with these components showed remarkable correspondence with the established mechanism-based categories of persistent pain, providing novel insight into the neurobiological signatures unique to nociceptive, neuropathic, and nociplastic pain.

When interpreting these dimensions, it is crucial to note that a negative correlation does not imply an absence of a feature but rather a stronger association with the negatively correlated feature. Thus, we choose to discuss these dimensions in terms of spectra with distinct phenotypes at either end (A and B) representative of the negative and positive weights, respectively.

#### 2.2.1 Nociplastic spectrum

The first brain-body dimension represents widespread bodily symptoms, as shown in Figure 2. Phenotype 1A is associated with more pain, numbness, and pins and needles throughout all regions of the body. This phenotype is also associated with weaker connectivity within and between several brain networks. This largely involves the left and right S1 region of the postcentral gyrus. These results suggest that widespread bodily symptoms are associated with weaker connectivity between S1 and other regions associated with somatomotor and attention networks. This presentation is also correlated with weaker connectivity within the visual network, specifically the occipital cortex and fusiform gyrus, and within subcortical and limbic regions such as the parahippocampal gyrus, hippocampus and basal ganglia.

**Fig. 2.**
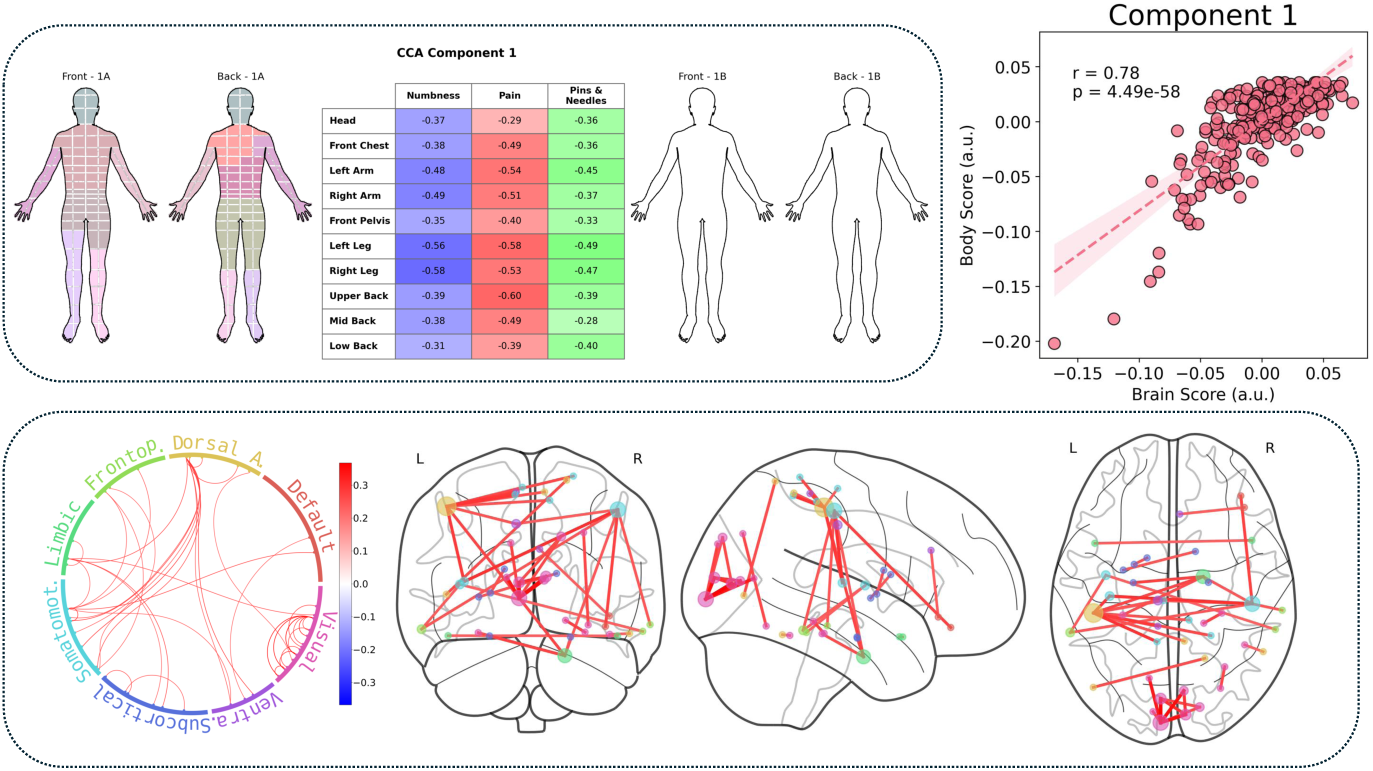
Canonical Component 1: The Nociplastic Dimension.

#### 2.2.2 Nociceptive-neuropathic spectrum

The second dimension represents a gradient from back pain on one end to numbness in the low back and limbs on the other, as shown in Figure 3. Phenotype 2B, the numbness phenotype, is associated with stronger connectivity between subcortical regions, specifically the thalamus and basal ganglia, and parts of the DMN, dorsal attention, and frontoparietal networks, while the converse is true for Phenotype 2A, the axial low back pain phenotype. This dimension is also associated with connectivity between the somatomotor and visual networks. Specifically, Phenotype 2B exhibits stronger connectivity between the primary somatosensory/motor cortices and regions of the right lateral occipital lobe.

**Fig. 3.**
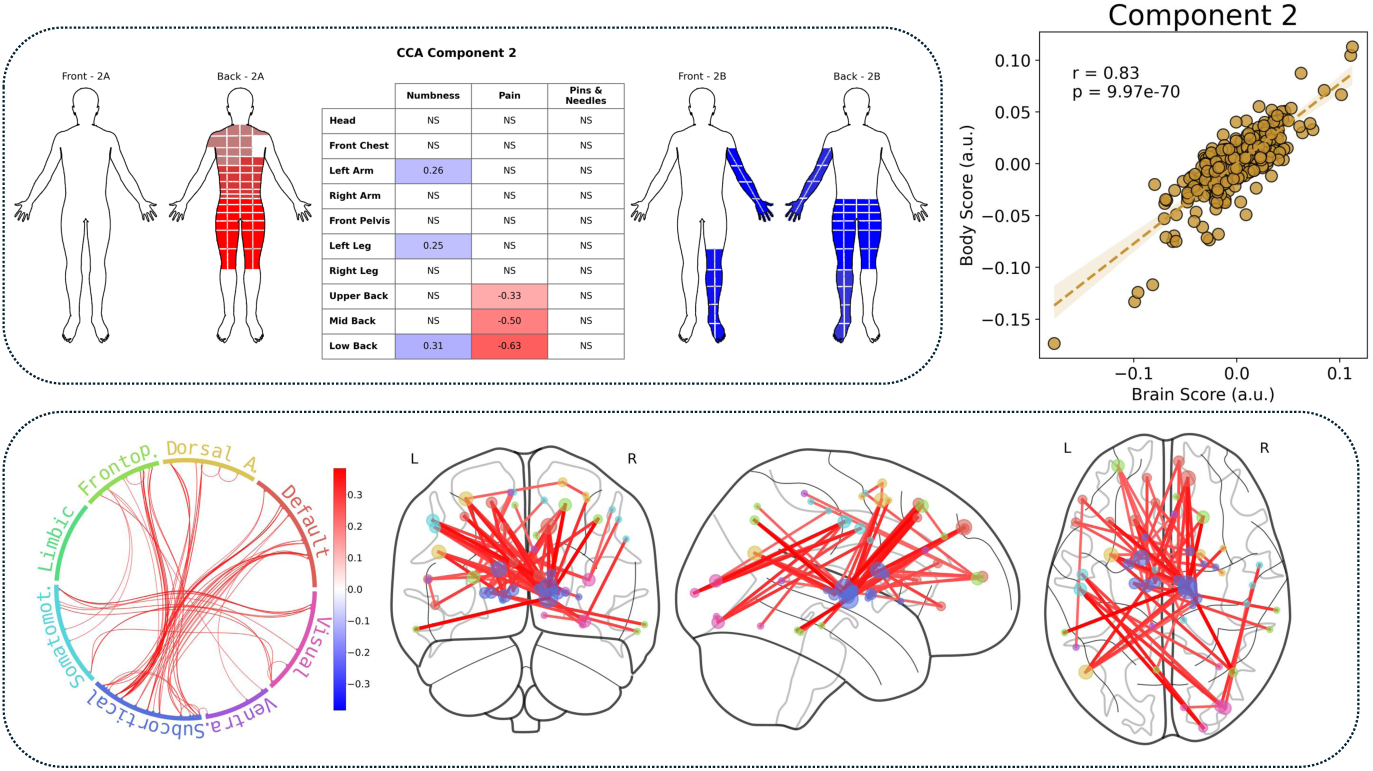
Canonical Component 2: Nociceptive-Neuropathic Dimension.

### 2.3 Model predicts spatial pain distribution in novel dataset

To test the generality of our model, we investigated whether it could predict body maps from resting-state data in an unseen dataset. Body maps were predicted for four novel subjects by projecting each subject’s resting-state data into the trained CCA space following the predictive framework outlined by Bilenko and Gallant Bilenko and Gallant (2016). In other words, an estimate for each body part and sensation was computed for each subject as a linear combination of his or her functional connectivity features and canonical weights calculated on our LBP cohort. Results of this test are shown in Figure 4. Visual inspection of these results indicates moderate correlation between predicted and true values, while a preliminary quantitative analysis suggests moderate predictive performance (accuracy score range 0.5-0.9).

**Fig. 4.**
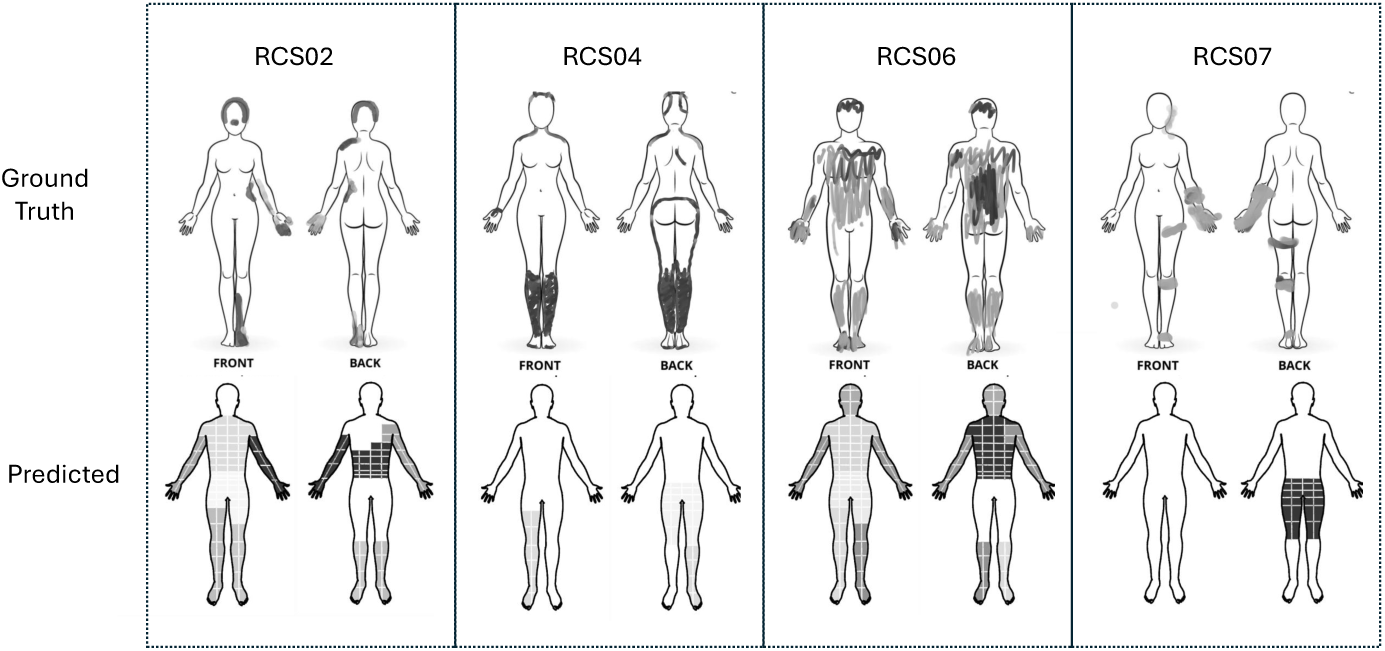
Pain body map predictions on novel dataset. Upper row is ground truth, lower row is predicted. All images are shown in grayscale to enable interpretation.

## 3 Discussion

With this study, we sought to identify neurobiological signatures underlying different phenotypes in our cLBP cohort. Our cross-decomposition analysis yielded seven brain-body dimensions, each representing a novel spectrum of somatosensory patterns correlated with resting-state functional connectivity. These dimensions form a low-dimensional space capable of predicting pain body maps in a previously unseen dataset, suggesting that these relationships extend beyond cLBP to chronic pain conditions more broadly.

Our model identified a subset of resting-state functional connectivity features associated with somatosensory patterns. This involved many brain areas traditionally associated with nociceptive processing, including S1, thalamus, and frontal regions, as well as some less expected regions, such as the inferior temporal gyrus and lateral occipital cortex. The wide range of regions and networks in our predictive set is consistent with previous reports associating chronic low back pain with global brain network reorganization Mansour et al (2016).

Our multivariate approach represents a departure from the discrete categorization normally employed in studies of chronic pain. Rather than identifying subjects as belonging to a specific pain type, we show that patients can be distributed along a set of spectra. We also want to highlight the relative simplicity of this model. The continuous linear distributions shown for each dimension indicate not just that brain and body patterns are associated but that they are directly linearly correlated, suggesting that an incremental change in one set of features may predict change in another.

Our first dimension shows an association between widespread bodily symptoms and weaker connectivity between a network of brain regions with hubs in bilateral S1. This phenotype is also associated with cognitive dysfunction, specifically elevated anxiety and fatigue and diminished social functioning (Supplementary Figure D9). This aligns with the emerging definition of nociplastic pain and provides a novel view into the related neurobiological mechanisms. S1 abnormalities, such as increased inflammatory activation, are increasingly recognized as a potential biomarker of nociplastic pain Shraim et al (2024). The involvement of occipital lobe interconnectivity is less expected. However, we are not the first to note visual network abnormalities in chronic pain subjects Shen et al (2019). We hypothesize that this may be indicative of widespread sensory circuit reorganization.

We refer to our second dimension as a nociceptive-neuropathic spectrum. We show that the amount of self-reported numbness in the back and legs (characteristic of neuropathic phenotypes) is associated with stronger connectivity between the thalamus and various regions of the cortex associated with attention and higher cognitive functions, as well as between somatomotor regions and parts of the occipital cortex. The strong association with thalamic connectivity is consistent with previous literature suggesting altered thalamocortical activity as a signature of neuropathic pain Garcia-Larrea and Peyron (2013); Alshelh et al (2016). Additionally, higher levels of cortical disinhibition have been shown in neuropathic pain patients compared to nociceptive patients Schwenkreis et al (2010). It is possible that the somatomotor-visual connectivity seen in our neuropathic phenotype represents an extension of this effect observable at rest.

Perhaps the most compelling aspect of our findings is the ability to predict pain body maps in an independent dataset. Despite the numerous potential sources of variation in the novel data, including out-of-distribution subjects and differences in data acquisition protocols, our model demonstrated moderate ability to predict spatial locations and intensity of pain from resting-state fMRI data. This is especially noteworthy given the relative simplicity of our model, employing only a weighted linear combination of a relatively small subset of brain connectivity features. While we recognize the limitations of this very small and largely qualitative example, we intend to further explore this prospect in future work.

Our work has several limitations. Primarily, we want to acknowledge the exploratory nature of this study; CCA analysis can only show correlations but does not provide any insight into causal links between our datasets, precluding any direct clinical utility. Additionally, our findings are restricted to individuals with cLBP who already exhibit altered neurocircuitry; this model is unlikely to generalize to healthy controls and may not even extend to other chronic pain conditions, although our preliminary results suggest a degree of universality. Methodologically, the subjective nature of body map reporting presents challenges in standardization and processing that could influence our results. Furthermore, we did not exclude participants based on medication use aside from opioids, which may confound the observed brain-body relationships.

Despite these limitations, the strong correlations and significance observed in our model suggest that it does capture meaningful associations between somatosensory abnormalities and brain connectivity in our cLBP cohort. These findings present resting-state functional connectivity as a potential avenue for deriving pain biomarkers. Additionally, the resemblance of our components to documented chronic pain phenotypes further supports the utility of our approach for uncovering latent neurobiological signatures in the chronic pain population that may be leveraged to provide more targeted pain treatments.

## 4 Methods

### 4.1 Participants

The data used in this analysis were collected as part of the ongoing comeBACK study (NIH Back Pain Consortium-BACPAC, 1U19AR076737-01), a multisite longitudinal study of chronic lower back pain Hue et al (2024). All participants are adults age ≥ 18 who had experienced low back pain for at least three months and at least half the days of the past six months at the time of recruitment and who reported that the low back was their dominant pain location. Inclusion criteria were developed with the intention to enroll non-specific cLBP patients, meaning that participants with a history of disorders such as cancer, spine infection, vertebral fracture, and several autoimmune diseases were excluded. Other exclusion criteria included common contraindications for MRI (claustrophobia, internal hardware) and an inability to complete all time points Hue et al (2024).

From an initial cohort of 449 cLBP patients, we selected subjects for whom body maps and baseline fMRI were available and who were not taking opioid medication at the time of data collection. This resulted in a dataset of 275 individuals (117 males and 158 females, age: 54 ± 16, mean ± SD, age range: 18–91 years old).

### 4.2 Body maps acquisition

Participants filled out body maps on paper in front of an examiner during the baseline visit. They received blank diagrams of the front and back of a body and were instructed to shade in all areas where they had felt each of three sensations (pain, numbness, and pins and needles) over the past 7 days.

### 4.3 Body maps processing

To convert the body maps data into a vector input suitable for CCA analysis, we aimed to extract a low-dimensional set of somatosensory patterns with which all subjects’ data could be described. To this end, a processing pipeline was designed with the intention of eliminating noise (e.g. inherent subject differences in marking style) while retaining valuable signal (e.g. a larger mark may indicate more salient pain). This pipeline is depicted in Figure 5, and additional details are provided in Appendix A.

**Fig. 5.**
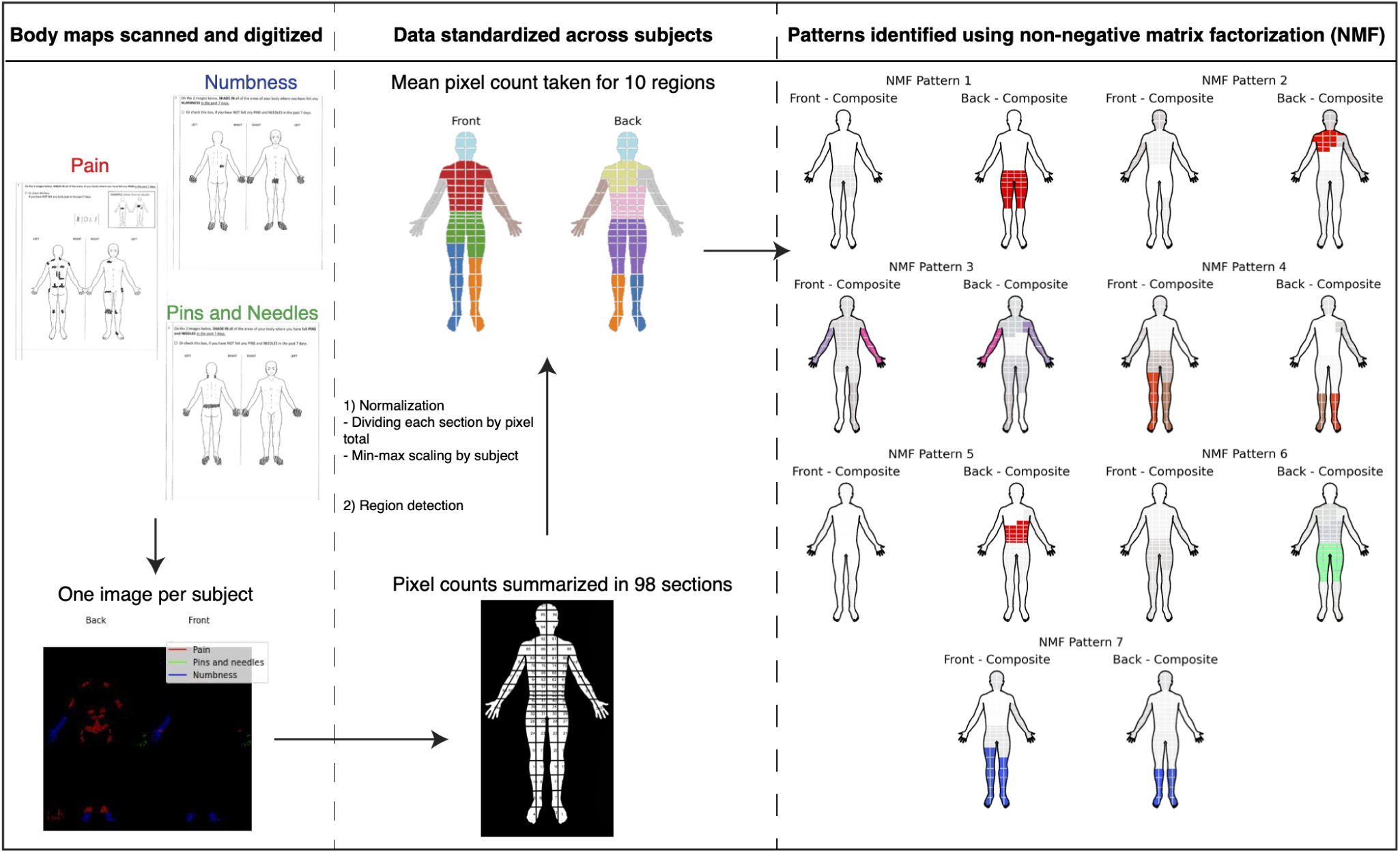
Body maps processing pipeline.

For each of the three sensations (pain, numbness, and pins and needles), raw pixel counts were summed within 98 grid sections on the front and back, resulting in 196 grid section values for each sensation per subject. A two-step normalization procedure was applied to the raw pixel count data to achieve comparable values across all grid sections and subjects. This involved first dividing each section’s pixel count data by the total number of pixels in that section (i.e., column normalization) and then scaling each subject’s data between 0 and 1 (i.e., row normalization).

Next, in order to increase SNR and further account for inter-subject variability, we aimed to summarize section values within meaningful anatomical regions. A data-driven approach was used to assign grid sections to larger anatomical regions based on group-level similarity in pixel count patterns while accounting for spatial proximity. Spatially-aware clustering was performed using a method from graph theory called community detection. First, a graph (similarity matrix, *S*) was constructed as the covariance matrix (*Cov*) of all subjects’ pixel data weighted by a spatial adjacency matrix (*A*), as defined below:

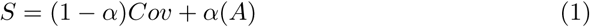

*A* was constructed such that sections that share a boundary in 2D (front or back grid) have edge value = 1 and mirrored front-back sections have edge value = 0.5. Communities were detected using the Louvain Community Detection Algorithm implemented in the python package *NetworkX* Hagberg et al (2008). Values of *α* were swept through 0-1 and the value that optimized modularity was chosen. The algorithm was then rerun using the optimal *α* value, and the mean normalized pixel count percentage was taken for all sections within each community (body part) for all three sensations.

Finally, nonnegative matrix factorization (NMF) was performed on these data to identify a sparse set of body map factors, referred to as “body map patterns” or simply “patterns”. The optimal number of NMF factors was determined using 5-fold cross-validation over 10 iterations with a range of factors from 2 to 10, selecting the number that minimized the acceleration of the cross-validation error curve (e.g. the elbow method). Descriptive statistics for each of the patterns are provided as the percent weight applied to each feature in the input “body part-sensation” space, with feature weights constituting more than 5% reported. Subjects’ values for all of the resulting body map patterns were used as input to the rCCA algorithm.

### 4.4 MRI data acquisition

Neuroimaging data were collected at four sites: University of California, San Francisco (UCSF), University of California, Davis (UCD), University of California, San Diego (UCSD), University of California, Irvine (UCI). All data were collected on 3T scanners from Siemens or GE. The protocol included structural T1-weighted scans and functional MRI with BOLD contrast with parameters as follows:

T1 structural scan: Repetition time (TR): 2500 ms (Siemens), 6 ms (GE); Echo time (TE): 3 ms; flip angle: 8; voxel size: 1 × 1 × 1 mm.

BOLD fMRI: TR: 800 ms; TE: 30 ms; flip angle: 52; voxel size: 2.4 x 2.4 x 2.4 mm.

### 4.5 fMRI preprocessing

Neuroimaging data were preprocessed using *fMRIPrep* 20.2.7 Esteban et al (2018b,a) and *Nilearn* Abraham et al (2014). A full fMRIPrep boilerplate file is provided in the Extended Data (Appendix E). Briefly, skull-stripped T1-weighted (T1w) MP-RAGE images were used for brain tissue segmentation and registration into standard MNI space. For BOLD data, susceptibility distortion correction was applied using fMRIPrep’s fieldmapless approach. The BOLD reference was co-registered to the T1w reference using boundary-based registration with 9 degrees of freedom. Head motion parameters were estimated before spatiotemporal filtering. BOLD runs were slice-time corrected and resampled into their original space and standard spaces (MNI152NLin2009cAsym, MNI152NLin6Asym). Several confounding time-series were calculated, including framewise displacement, DVARS, and global signals.

Further denoising and time series extraction was performed using the *NiftiLabelsMasker* class in Nilearn. This included spatial smoothing (5-mm FWHM Gaussian kernel), temporal bandpass filtering (high pass = discrete cosines transformation, low pass = 0.08 Hz), nuisance signal regression (24 motion parameters, 2 white matter/CSF parameters), and scrubbing with a framewise displacement threshold of 0.5 mm. Subjects with less than 5 minutes of data retained after motion scrubbing were excluded from analysis. The Brainnetome atlas Fan et al (2016) was applied to extract mean BOLD time signals from 246 anatomically and functionally defined regions. Pearson’s correlation was applied to each subject’s time series to obtain a 246 x 246 functional correlation matrix (“functional connectome”/ FC). Each subject’s FC was standardized using Fishers r-to-z transform.

### 4.6 Functional Connectivity Feature Selection

Feature selection was performed to identify a subset of the 30135 unique FC edges to use in CCA analysis using a resampled feature selection following the method described in Buch et al., 2023 Buch et al (2023). Over 100 iterations, Spearman’s rank correlation coefficient was calculated between each FC edge and each body map pattern, and edges were ranked by the number of times they were found statistically correlated (*p <* 0.001) with at least one body map pattern.

### 4.7 Regularized CCA

Regularized CCA (rCCA) was performed between the body map patterns (Y) and FC features (X) using the python package *pyrcca* Bilenko and Gallant (2016). Parameter optimization was conducted through a repeated grid search over three variables: the regularization parameter *λ*, the number of canonical components (*N_CC_*), and the number of FC features (*N_F C_*). For each grid search iteration, 99% of the dataset was randomly selected to train pyrcca’s *CCACrossValidate* object class, which performs K-fold cross-validation to optimize *λ* and *N_CC_*. This was performed using a variable number of top-ranked FC features ranging from 100 to 400, selecting the hyperparameter triplet that maximized the mean correlation over all components.

The number of canonical components calculated in CCA may be less than or equal to the minimum number of features in the input matrices. Given a lack of *a priori* knowledge regarding the optimal number of latent brain-body dimensions, an initial course grid search was performed to optimize the number of canonical components used in the model. Over 100 iterations, the grid search procedure described above was performed using 10-fold cross-validation with a hold-out size of 20% and a parameter grid as follows: *λ*: [10*^−^*^7^ to 10^4^, 20 logarithmically spaced values], *N_CC_*: [1 to 7 (the number of body map patterns)], *N_F C_*: [100 to 400, step size of 20]. The most frequently occurring number of components over all grid search iterations was selected for the final model.

A bootstrapped permutation test was then performed to evaluate reproducibility in held-out data and to establish significance of the resulting canonical components (brain-body dimensions). Over 1000 iterations, data were randomly split into a training set and test set (test set size = 1%, 28 subjects). Grid search with 10-fold cross-validation was performed on the training set to optimize *λ* and *N_F C_* while restricting *N_CC_* to the optimal number obtained previously, with a refined parameter grid of *λ*: [10*^−^*^8^ to 10^3^, 30 linearly spaced values] and *N_F C_*: [180 to 400, step size of 20]. An rCCA model was then trained on the whole training set using the optimized parameters, and the held-out test set was projected into the trained rCCA space using pyrcca’s *validate()* method. These steps were then repeated with the rows of Y randomly permuted to establish a null distribution, yielding a set of 1000 canonical correlations for both the true and null test datasets. These distributions were compared using Welch’s t-test testing the one-sided alternative hypothesis that null set correlations are greater. A violin plot of the true and null canonical correlations is shown in Appendix B.

Finally, rCCA was performed on the entire dataset. Parameter optimization was performed with 30-fold cross-validation using the same parameter grid, and a final rCCA model was trained using these optimized parameters, yielding a “brain-dimension” and “body-dimension” score for each subject and component. Spearman’s correlation was calculated between each FC feature and each brain dimension score, as well as between each body part-sensation value and each body dimension score. Significant correlations following Bonferroni correction are presented for each component (*p_brain_* = 0.00014, *p_body_* = 0.0016).

### 4.8 Association with demographic, clinical, and psychosocial risk factors

Post-hoc analysis was performed to investigate any associations between the final canonical components and common LBP risk factors. These were categorized by domain: demographic (biological sex, BMI, age), clinical (presence of imaging findings, score on Pain, Enjoyment and General Activity (PEG) scale, a 1-10 pain scale, mechanistic pain category), and psychosocial (Patient-Reported Outcomes Measurement Information System (PROMIS) measurements). A single score was obtained for each subject and component by projecting the subject’s location in each component space to the *y* = *x* line. Correlation was then calculated between each subject’s component score and each variable of interest. Spearman Correlation was used for continuous variables (scatter plots), Mann-Whitney U test was used for binary categorical variables, and Kruskal-Wallis test was used for categorical variables with more than 2 groups. Results are shown in the Extended Data (Appendix D; Supplementary Figures D7 - D9).

### 4.9 Details of novel dataset

Resting-state data and pain body maps were provided for four novel subjects collected as part of a research trial of deep brain stimulation for refractory chronic neuropathic pain. Patient demographics and body map acquisition protocols have been previously described Kwong et al (2023). Briefly, participants completed digital body maps indicating pain location and intensity multiple times a day over a ten-day inpatient hospital stay. Data used in this analysis were collected at baseline.

Profiles of each of the four participants included in this study are provided below:

- RCS02: 58 year old female with history of right insular stroke with post stroke pain syndrome
- RCS04: 54 year old female with spinal degenerative joint disease and chemotherapy induced neuropathy in feet and ankles
- RCS06: 49 year old male with cervical spine stenosis and spinal cord injury at T4
- RCS07: 51 year old female with pontine hemorrhage and One-and-a-half syndrome

Functional connectomes were derived for each novel subject using the same brain atlas and neuroimaging processing protocol described above. Predicted body map features were then computed as the dot product between the inverse of the body map weights and the weighted novel functional data:

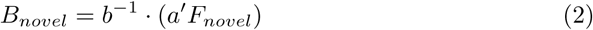

where *a* and *b* are the canonical weight vectors for the brain and body axes, respectively.

Model accuracy was assessed using binarized pain values for each body part. The presence or absence of pain was visually evaluated for each of the ten body parts per subject in the true data. These values were compared against binarized predicted body maps to generate an accuracy score per subject, i.e. the ratio of body parts with matching labels in true and predicted data.

## Acknowledgements

Research reported in this publication was supported by the National Institute Of Arthritis And Musculoskeletal And Skin Diseases of the National Institutes of Health under Award Number U19AR076737. The content is solely the responsibility of the authors and does not necessarily represent the official views of the National Institutes of Health.

## Declarations

### 4.11 Data availability

The datasets generated during and/or analysed during the current study will be made available on the Vivli Platform prior to publication, [https://search.vivli.org/studyDetails/fromSearch/7a7b72f2-7086-4344-89ad-957c641c8c6c].

### 4.12 Author contribution

J.A.C conceived and conducted main analyses and wrote paper. A.B. digitized and performed preliminary analysis on body map data. J.M. and P.S. provided novel dataset. J.M. processed fMRI data in novel dataset. All authors provided critical feedback, reviewed, and edited the paper.

## Appendix A Body map dimensionality reduction

Community detection applied to the weighted covariance matrix of gridded pixel count data yielded a solution of 10 communities corresponding to distinct body parts, with an optimized weighting factor *α* = 0.90 and a peak modularity = 0.71. These body parts are illustrated in Figure 5. Five of these communities encompassed grid sections from both the front and back body maps (head, left arm, right arm, left leg, and right leg) while five were restricted to either the front body (front chest, front pelvis) or back body (upper back, mid back, low back).

For each sensation, the mean normalized pixel count was calculated across all sections within each body part, yielding a vector of size 30 (number of sensations x number of body parts) for each subject. This matrix was then used as input to the NMF algorithm. The optimal number of factors was empirically selected to be 7 and confirmed with visual inspection of the cross-validation error curve. The resulting patterns are represented in Figure 5 of the main text.

The NMF reconstruction explained 79.60% of the variance of the input data. Pattern 1 explained a majority of the variance (29.25%) and represents low back pain predominantly (88.37% feature weight) with some front pelvis pain (7.53%). All other patterns each contributed an additional 6 to 12% variance. Pattern 2 represents upper body pain, mostly upper back pain (58.22%) as well as some right arm pain (11.48%) and head pain (10.39%). Pattern 3 broadly represents arm symptoms, including left arm pain (22.05%), numbness (14.15%), and pins and needles (5.70%) as well as right arm numbness (11.34%) and pain (8.05%). Pattern 4 encompasses leg symptoms, including right leg pain (34.47%); pins and needles (9.11%) and left leg pain (22.17%); pins and needles (6.69%) as well as some front pelvis pain (8.60%) and right arm pain (5.56%). Pattern 5 is predominantly mid back pain (75.63%). Pattern 6 represents numbness and pins and needles throughout the back, with weights as follows: low back pins and needles (34.75%), low back numbness (21.92%), mid back numbness (6.75%), upper back pins and needles (5.16%), front pelvis pain (5.03%). Finally, Pattern 7 represents more leg symptoms, including left leg numbness (32.15%), pins and needles (10.61%) and pain (5.65%), and right leg numbness (25.13%) and pins and needles (5.26%).

**Fig. B1.**
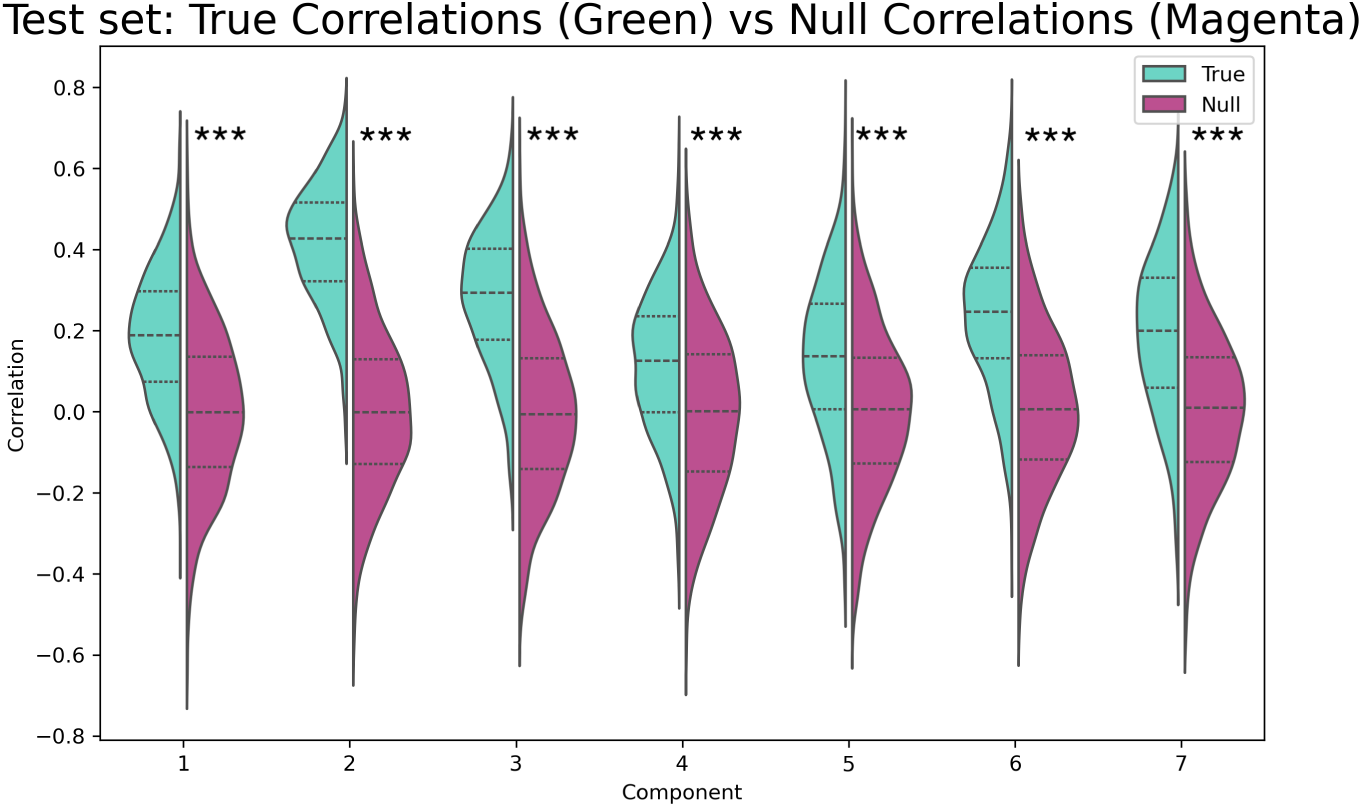
Correlation of CCA components in true data vs. permuted data.

## Appendix B Permutation test for significance

Following the initial grid search, a 7-component solution was selected for further analysis, with an optimized *N_CC_* = 7 in 40% of the grid search iterations. All 7 dimensions were significant in held-out test sets compared to shuffled data (Component 1: Mean *r* = 0.19, *p* = 3.29e-110; Component 2: Mean *r* = 0.41, *p* = 0; Component 3: Mean *r* = 0.28, *p* = 2.31e-215; Component 4: Mean *r* = 0.12, *p* = 4.84e-48; Component 5: Mean *r* = 0.13, *p* = 2.83e-47; Component 6: Mean *r* = 0.24, *p* = 4.17e-145; Component 7: Mean *r* = 0.19, *p* = 4.49e-95). Violin plots showing true and null correlations are presented in Figure B1.

**Fig. C2.**
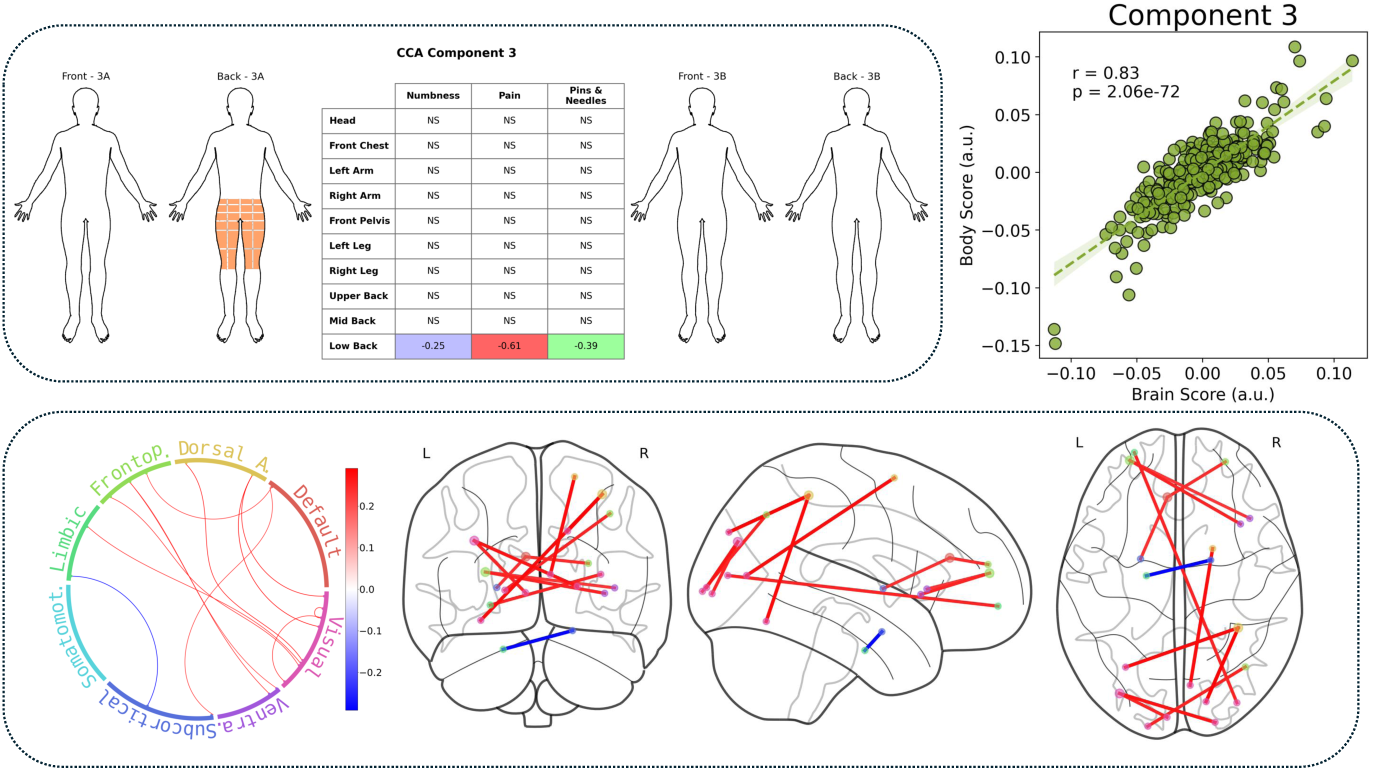
Brain and body feature weights for Component 3.

## Appendix C Additional Canonical Component Weights

Phenotype 3A (Figure C2) represents low back symptoms, with low back pain, numbness, and pins and needles all negatively correlated with this component. This phenotype shows stronger connectivity between the amygdala and parahippocampal gyrus, and weaker cortico-cortical connectivity between regions throughout the frontal, parietal, occipital, and cingulate cortices and the insula, as well as between the basal ganglia and the cingulate.

The fourth dimension (Figure C3) represents a superior-inferior gradient, with phenotype 4A showing upper body symptoms, including pain and numbness in the arms and upper to mid back, and 4B showing low body symptoms, such as low back pain and leg numbness and pins and needles. Phenotype 4A is associated with stronger connectivity between somatomotor regions, including S1, and DMN, attention, and limbic regions throughout the cortex. Phenotype 4B shows stronger connectivity between the thalamus and superior frontal gyrus.

Phenotype 5A is associated with pain in the legs and pelvis, while 5B involves pain in the upper left back and numbness in the left leg (Figure C4). Phenotype 5A shows stronger connectivity within and between various somatomotor, attention, and default regions, as well as between the thalamus and parts of the occipital cortex. Phenotype 5B shows stronger interhemispheric connectivity in the superior temporal gyrus.

Phenotype 6A is associated with low back pain, while 6B shows pins and needles in the upper and lower back and pain in the mid back (Figure C5). Phenotype 6A shows stronger connectivity between visual regions, in particular the left and right lateral occipital cortex, as well as the cingulate and precuneus. Phenotype 6B shows stronger connectivity between the superior parietal lobule and parts of the lateral occipital cortex.

**Fig. C3.**
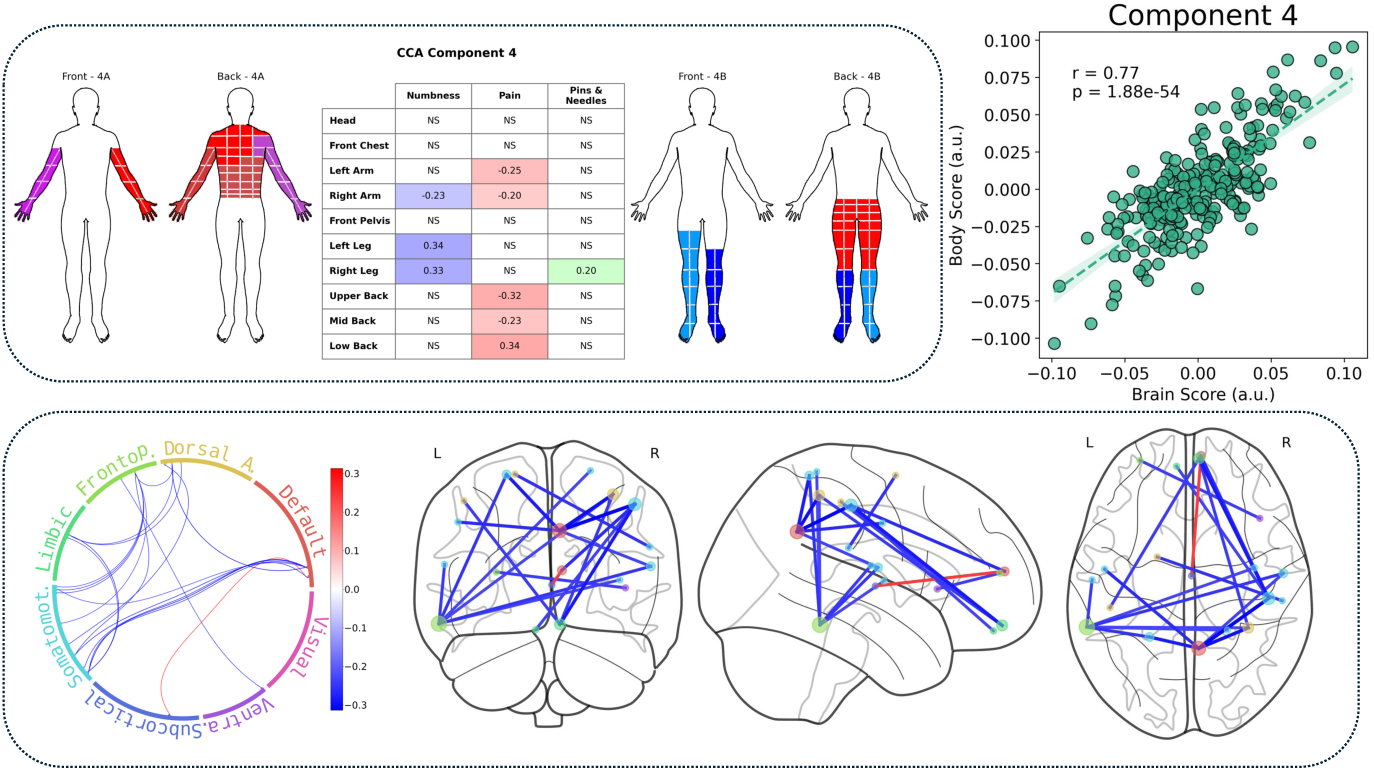
Brain and body feature weights for Component 4.

**Fig. C4.**
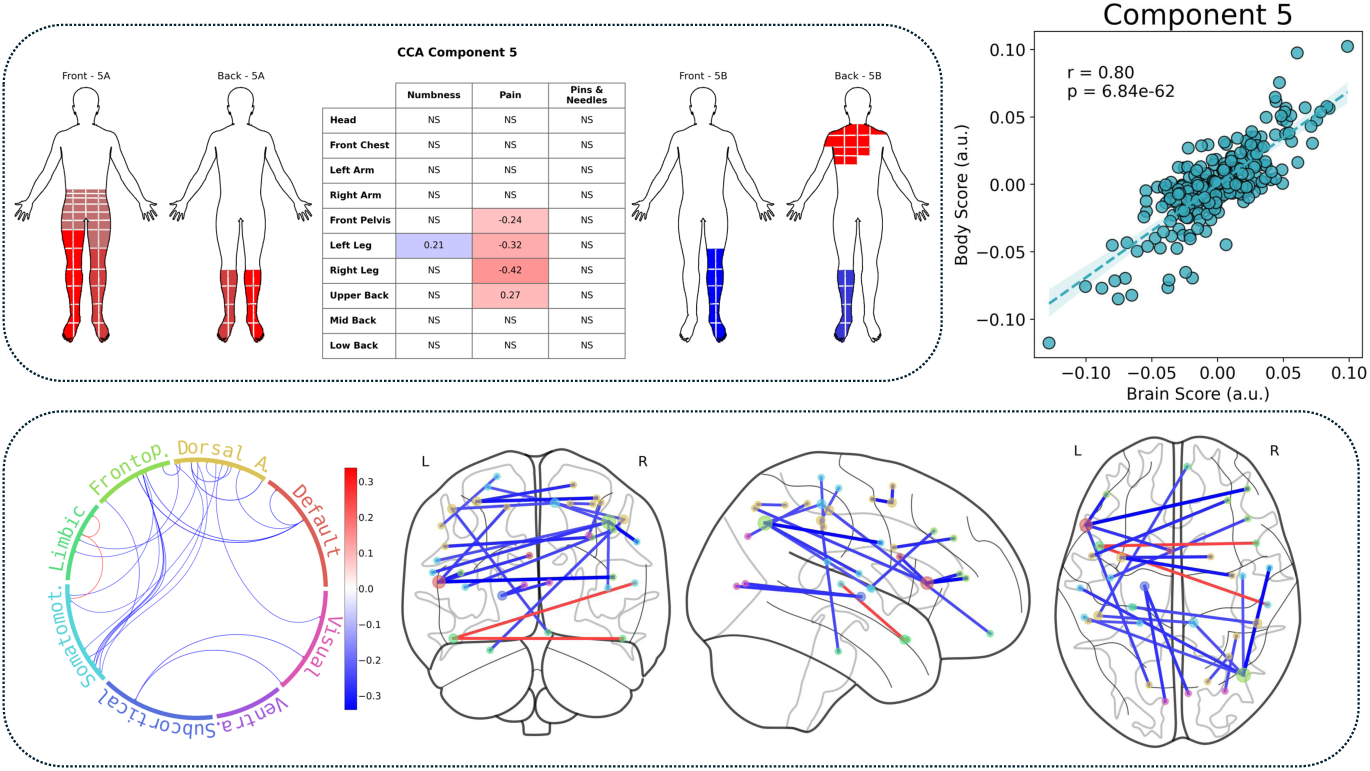
Brain and body feature weights for Component 5.

Component 7 shows upper back pain on one end and mid back pain on the other. Phenotype 7B, the mid back pain phenotype, shows stronger connectivity between the left inferior frontal gyrus and the left insula, cingulate, and parahippocampal gyrus, as well as within the right parahippocampal gyrus.

**Fig. C5.**
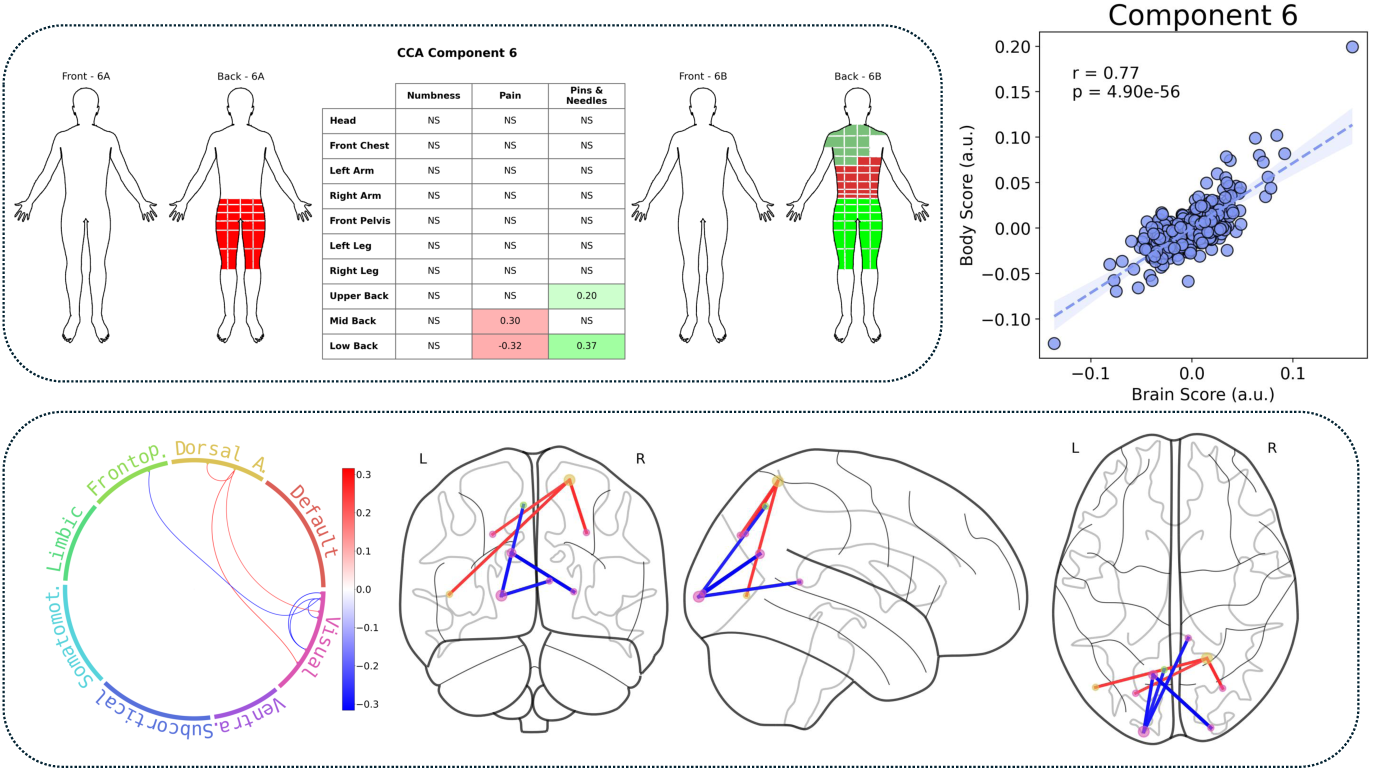
Brain and body feature weights for Component 6.

**Fig. C6.**
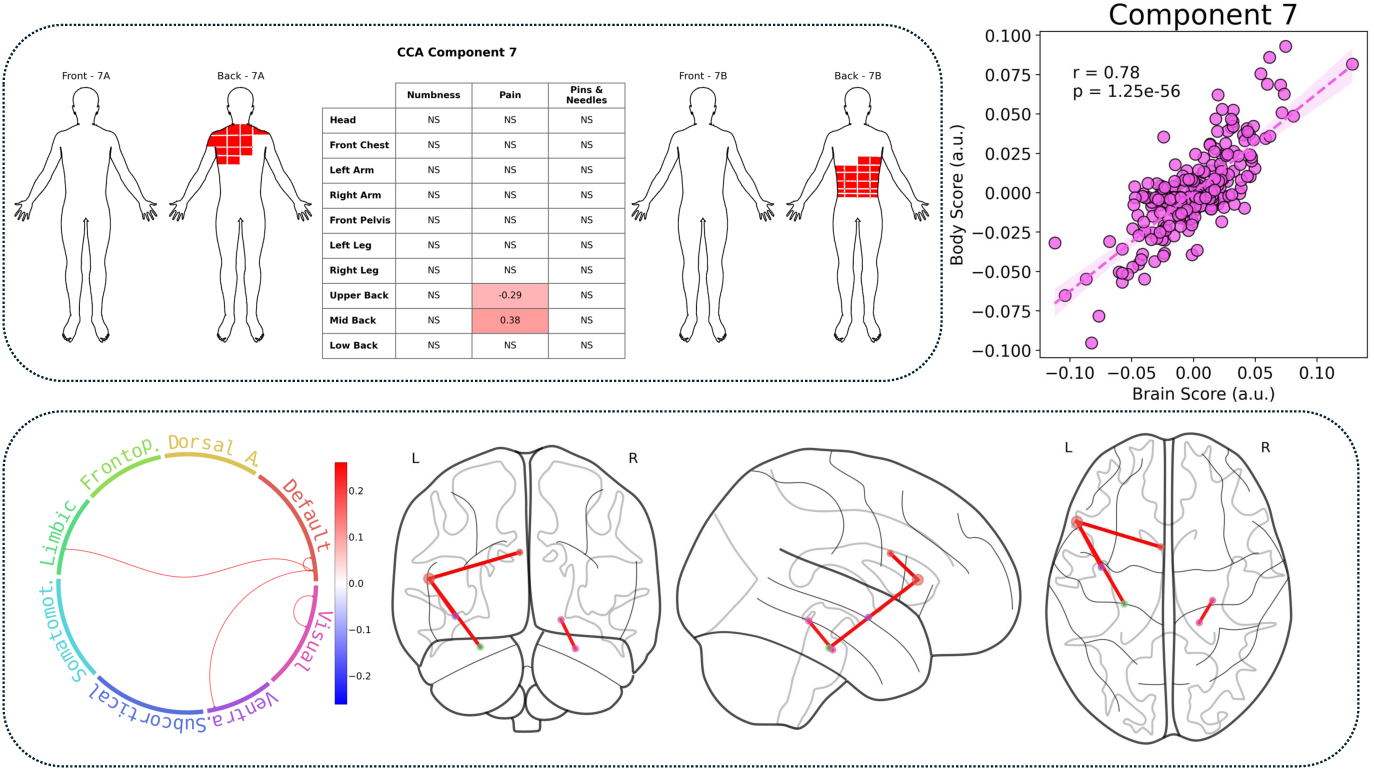
Brain and body feature weights for Component 7.

## Appendix D Association with biopsychosocial factors

We investigated associations between each component score and a set of biopsychosocial risk and prognostic factors for LBP. Results are shown in Figures D7 - D9. Each column represents a component, and each row is a variable. Tests with *p* ≤ 0.05 are indicated in the subplot title.

Component 1 is associated with almost all PROMIS variables with the exception of depression, suggesting that a tendency to report more bodily symptoms is also correlated with higher scores on negatively valanced PROMIS indicators (anxiety, fatigue, sleep disturbance) and lower scores on positively valanced PROMIS indicators (social roles and activities, cognitive function, physical function). Phenotype 1A is also associated with a higher PEG score and a higher prevalence of Modic Type 1 imaging findings. Component 2 showed an association with sex, with females tending more toward Phenotype 2B. Phenotype 3A is correlated with lower age and a higher score on the PROMIS fatigue scale. Component 7 is associated with several demographic factors, with Phenotype 7B correlated with female sex, higher BMI, and lower age. It is also associated with higher rates of social activity.

**Fig. D7.**
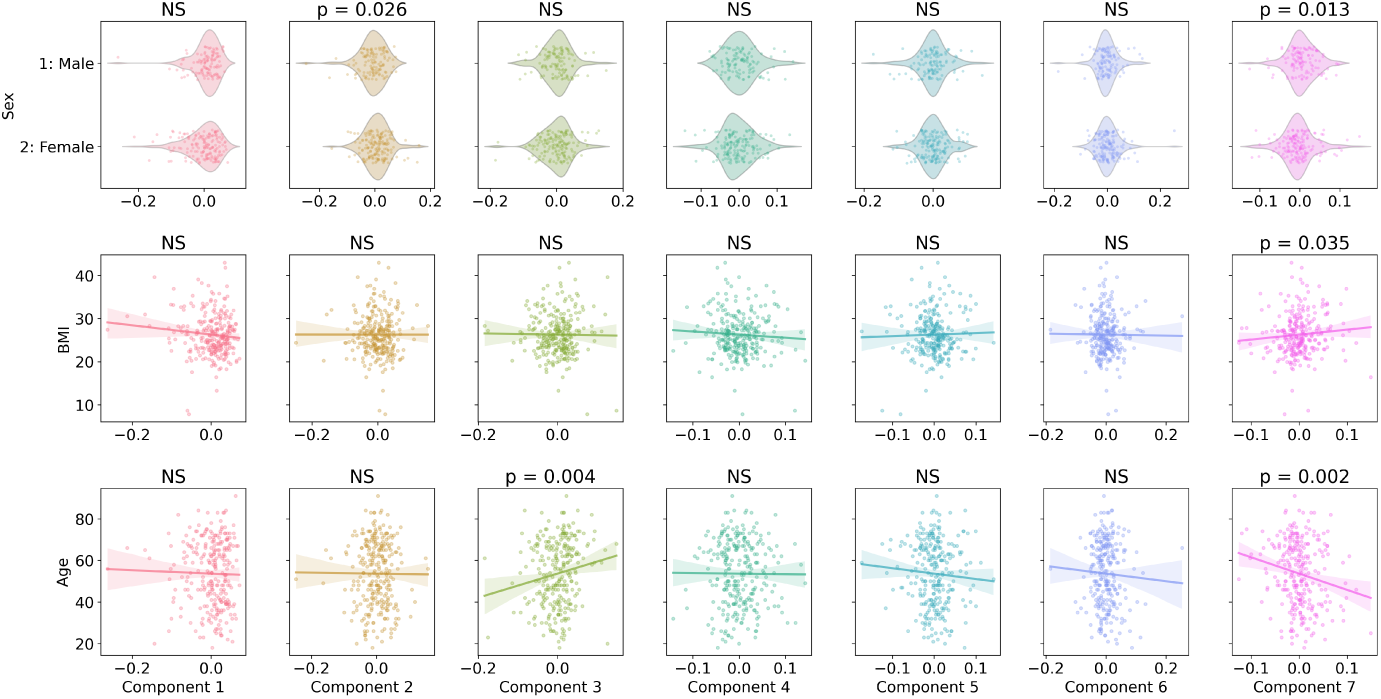
Demographic Variables.

**Fig. D8.**
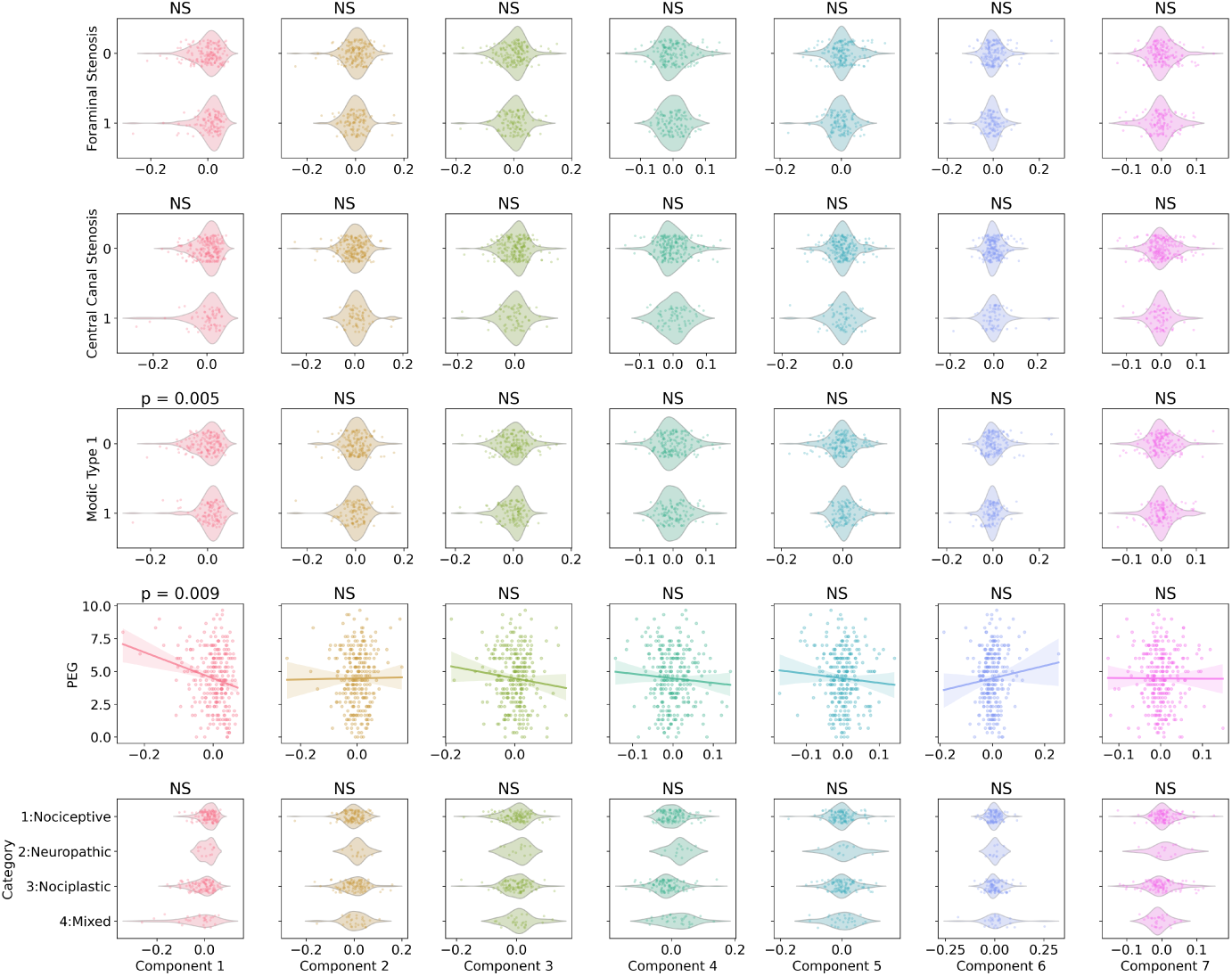
Clinical Variables.

## Appendix E fMRIPrep boilerplate text

Results included in this manuscript come from preprocessing performed using *fMRIPrep* 20.2.7 (Esteban et al (2018b); Esteban et al (2018a); RRID:SCR 016216), which is based on *Nipype* 1.7.0 (Gorgolewski et al (2011); Gorgolewski et al (2018); RRID:SCR 002502).

### E.1 Anatomical data preprocessing

A total of 1 T1-weighted (T1w) images were found within the input BIDS dataset. The T1-weighted (T1w) image was corrected for intensity non-uniformity (INU) with N4BiasFieldCorrection Tustison et al (2010), distributed with ANTs 2.3.3 (Avants et al, 2008, RRID:SCR 004757), and used as T1w-reference throughout the workflow. The T1w-reference was then skull-stripped with a *Nipype* implementation of the antsBrainExtraction.sh workflow (from ANTs), using OASIS30ANTs as target template. Brain tissue segmentation of cerebrospinal fluid (CSF), white-matter (WM) and gray-matter (GM) was performed on the brain-extracted T1w using fast (FSL 5.0.9, RRID:SCR 002823, Zhang et al, 2001). Brain surfaces were reconstructed using recon-all (FreeSurfer 6.0.1, RRID:SCR 001847, Dale et al, 1999), and the brain mask estimated previously was refined with a custom variation of the method to reconcile ANTs-derived and FreeSurfer-derived segmentations of the cortical gray-matter of Mindboggle (RRID:SCR 002438, Klein et al, 2017). Volume-based spatial normalization to two standard spaces (MNI152NLin2009cAsym, MNI152NLin6Asym) was performed through nonlinear registration with antsRegistration (ANTs 2.3.3), using brain-extracted versions of both T1w reference and the T1w template. The following templates were selected for spatial normalization: *ICBM 152 Nonlinear Asymmetrical template version 2009c* Fonov et al (2009), RRID:SCR 008796; TemplateFlow ID: MNI152NLin2009cAsym, *FSL’s MNI ICBM 152 non-linear 6^th^Generation Asymmetric Average Brain Stereotaxic Registration Model* Evans et al (2012), RRID:SCR 002823; TemplateFlow ID: MNI152NLin6Asym.

**Fig. D9.**
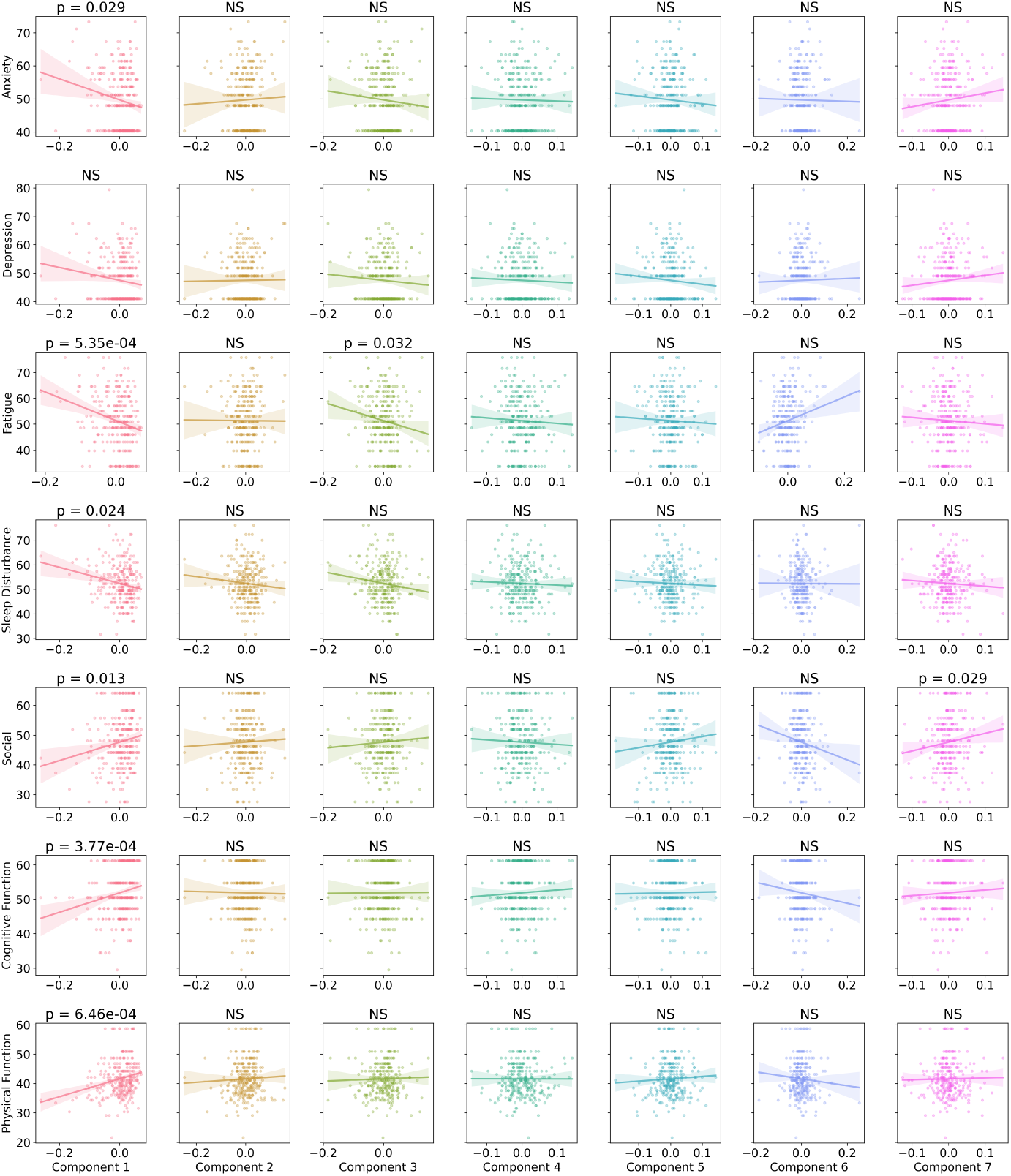
Psychosocial Variables. Anxiety: PROMIS Anxiety 4a (higher = more anxiety); Depression: PROMIS Depression 4a (higher = more depressed); Fatigue: PROMIS Fatigue 4a (higher = more fatigued); Sleep Disturbance: PROMIS Sleep Disturbance (higher = more disturbance); Social: PROMIS Social Roles and Activities 4a (higher = more social); Cognitive Function: PROMIS Cognitive Function 2a (higher = more cognitive function), Physical Function: PROMIS Physical Function 6b (higher = more function).

### E.2 Functional data preprocessing

For each of the 1 BOLD runs found per subject (across all tasks and sessions), the following preprocessing was performed. First, a reference volume and its skull-stripped version were generated by aligning and averaging 1 single-band references (SBRefs). A deformation field to correct for susceptibility distortions was estimated based on *fMRIPrep*’s *fieldmap-less* approach. The deformation field is that resulting from co-registering the BOLD reference to the same-subject T1w-reference with its intensity inverted Wang et al (2017); Huntenburg (2014). Registration is performed with antsRegistration (ANTs 2.3.3), and the process regularized by constraining deformation to be nonzero only along the phase-encoding direction, and modulated with an average fieldmap template Treiber et al (2016). Based on the estimated susceptibility distortion, a corrected EPI (echo-planar imaging) reference was calculated for a more accurate co-registration with the anatomical reference. The BOLD reference was then co-registered to the T1w reference using bbregister (FreeSurfer) which implements boundary-based registration Greve and Fischl (2009). Co-registration was configured with nine degrees of freedom to account for distortions remaining in the BOLD reference. Head-motion parameters with respect to the BOLD reference (transformation matrices, and six corresponding rotation and translation parameters) are estimated before any spatiotemporal filtering using mcflirt (FSL 5.0.9, Jenkinson et al, 2002). BOLD runs were slice-time corrected to 0.351s (0.5 of slice acquisition range 0s-0.703s) using 3dTshift from AFNI 20160207 (Cox and Hyde, 1997, RRID:SCR 005927). First, a reference volume and its skull-stripped version were generated using a custom methodology of *fMRIPrep*. The BOLD time-series (including slice-timing correction when applied) were resampled onto their original, native space by applying a single, composite transform to correct for head-motion and susceptibility distortions. These resampled BOLD time-series will be referred to as *preprocessed BOLD in original space*, or just *preprocessed BOLD*. The BOLD time-series were resampled into several standard spaces, correspondingly generating the following *spatially-normalized, preprocessed BOLD runs*: MNI152NLin2009cAsym, MNI152NLin6Asym. First, a reference volume and its skull-stripped version were generated using a custom methodology of *fMRIPrep*. Automatic removal of motion artifacts using independent component analysis (ICA-AROMA, Pruim et al, 2015) was performed on the *preprocessed BOLD on MNI space* time-series after removal of non-steady state volumes and spatial smoothing with an isotropic, Gaussian kernel of 6mm FWHM (full-width half-maximum). Corresponding “non-aggresively” denoised runs were produced after such smoothing. Additionally, the “aggressive” noise-regressors were collected and placed in the corresponding confounds file. Several confounding time-series were calculated based on the *preprocessed BOLD* : framewise displacement (FD), DVARS and three region-wise global signals. FD was computed using two formulations following Power (absolute sum of relative motions, Power et al (2014)) and Jenkinson (relative root mean square displacement between affines, Jenkinson et al (2002)). FD and DVARS are calculated for each functional run, both using their implementations in *Nipype* (following the definitions by Power et al, 2014). The three global signals are extracted within the CSF, the WM, and the whole-brain masks. Additionally, a set of physiological regressors were extracted to allow for component-based noise correction (*CompCor*, Behzadi et al, 2007). Principal components are estimated after high-pass filtering the *preprocessed BOLD* time-series (using a discrete cosine filter with 128s cut-off) for the two *CompCor* variants: temporal (tCompCor) and anatomical (aCompCor). tCompCor components are then calculated from the top 2% variable voxels within the brain mask. For aCompCor, three probabilistic masks (CSF, WM and combined CSF+WM) are generated in anatomical space. The implementation differs from that of Behzadi et al. in that instead of eroding the masks by 2 pixels on BOLD space, the aCompCor masks are subtracted a mask of pixels that likely contain a volume fraction of GM. This mask is obtained by dilating a GM mask extracted from the FreeSurfer’s *aseg* segmentation, and it ensures components are not extracted from voxels containing a minimal fraction of GM. Finally, these masks are resampled into BOLD space and binarized by thresholding at 0.99 (as in the original implementation). Components are also calculated separately within the WM and CSF masks. For each CompCor decomposition, the *k* components with the largest singular values are retained, such that the retained components’ time series are sufficient to explain 50 percent of variance across the nuisance mask (CSF, WM, combined, or temporal). The remaining components are dropped from consideration. The head-motion estimates calculated in the correction step were also placed within the corresponding confounds file. The confound time series derived from head motion estimates and global signals were expanded with the inclusion of temporal derivatives and quadratic terms for each Satterthwaite et al (2013). Frames that exceeded a threshold of 0.5 mm FD or 1.5 standardised DVARS were annotated as motion outliers. All resamplings can be performed with *a single interpolation step* by composing all the pertinent transformations (i.e. head-motion transform matrices, susceptibility distortion correction when available, and co-registrations to anatomical and output spaces). Gridded (volumetric) resamplings were performed using antsApplyTransforms (ANTs), configured with Lanczos interpolation to minimize the smoothing effects of other kernels Lanczos (1964). Non-gridded (surface) resamplings were performed using mri vol2surf (FreeSurfer).

## References

Abraham A, Pedregosa F, Eickenberg M, et al (2014) Machine learning for neuroimaging with scikit-learn. Frontiers in Neuroinformatics 8. 10.3389/fninf.2014.00014, URL https://www.frontiersin.org/articles/10.3389/fninf.2014.00014/full

Alshelh Z, Di Pietro F, Youssef AM, et al (2016) Chronic neuropathic pain: it’s about the rhythm. Journal of Neuroscience 36(3):1008–1018

Alshelh Z, Brusaferri L, Saha A, et al (2022) Neuroimmune signatures in chronic low back pain subtypes. Brain 145(3):1098–1110

Avants B, Epstein C, Grossman M, et al (2008) Symmetric diffeomorphic image registration with cross-correlation: Evaluating automated labeling of elderly and neurodegenerative brain. Medical Image Analysis 12(1):26–41. 10.1016/j.media.2007.06.004, URL http://www.sciencedirect.com/science/article/pii/S1361841507000606

Baliki MN, Petre B, Torbey S, et al (2012) Corticostriatal functional connectivity predicts transition to chronic back pain. Nature neuroscience 15(8):1117–1119

Behzadi Y, Restom K, Liau J, et al (2007) A component based noise correction method (CompCor) for BOLD and perfusion based fmri. NeuroImage 37(1):90–101. 10.1016/j.neuroimage.2007.04.042, URL http://www.sciencedirect.com/science/article/pii/S1053811907003837

Bilenko NY, Gallant JL (2016) Pyrcca: regularized kernel canonical correlation analysis in python and its applications to neuroimaging. Frontiers in neuroinformatics 10:49

van Boekel RL, Vissers KC, van der Sande R, et al (2017) Moving beyond pain scores: Multidimensional pain assessment is essential for adequate pain management after surgery. PLoS One 12(5):e0177345

Brinjikji W, Luetmer PH, Comstock B, et al (2015) Systematic literature review of imaging features of spinal degeneration in asymptomatic populations. American journal of neuroradiology 36(4):811–816

Brummett CM, Bakshi RR, Goesling J, et al (2016) Preliminary validation of the michigan body map. Pain 157(6):1205–1212

Buch AM, Vértes PE, Seidlitz J, et al (2023) Molecular and network-level mechanisms explaining individual differences in autism spectrum disorder. Nature neuroscience 26(4):650–663

Clauw DJ (2024) Why don’t we use a body map in every chronic pain patient yet? Pain 165(8):1660–1661

Cox RW, Hyde JS (1997) Software tools for analysis and visualization of fmri data. NMR in Biomedicine 10(4-5):171–178. 10.1002/(SICI)1099-1492(199706/08)10:4/5⟨171::AID-NBM453⟩3.0.CO;2-L

Dale AM, Fischl B, Sereno MI (1999) Cortical surface-based analysis: I. seg-mentation and surface reconstruction. NeuroImage 9(2):179–194. 10.1006/nimg.1998.0395, URL http://www.sciencedirect.com/science/article/pii/S1053811998903950

Drysdale AT, Grosenick L, Downar J, et al (2017) Resting-state connectivity biomarkers define neurophysiological subtypes of depression. Nature medicine 23(1):28–38

Esteban O, Blair R, Markiewicz CJ, et al (2018a) fmriprep. Software 10.5281/zenodo.852659

Esteban O, Markiewicz C, Blair RW, et al (2018b) fMRIPrep: a robust preprocessing pipeline for functional MRI. Nature Methods 10.1038/s41592-018-0235-4

Evans A, Janke A, Collins D, et al (2012) Brain templates and atlases. NeuroImage 62(2):911–922. 10.1016/j.neuroimage.2012.01.024

Fan L, Li H, Zhuo J, et al (2016) The human brainnetome atlas: a new brain atlas based on connectional architecture. Cerebral cortex 26(8):3508–3526

Fitzcharles MA, Cohen SP, Clauw DJ, et al (2021) Nociplastic pain: towards an understanding of prevalent pain conditions. The Lancet 397(10289):2098–2110

Fonov V, Evans A, McKinstry R, et al (2009) Unbiased nonlinear average age-appropriate brain templates from birth to adulthood. NeuroImage 47, Supplement 1:S102. 10.1016/S1053-8119(09)70884-5

Garcia-Larrea L, Peyron R (2013) Pain matrices and neuropathic pain matrices: a review. PAIN® 154:S29–S43

Gorgolewski K, Burns CD, Madison C, et al (2011) Nipype: a flexible, lightweight and extensible neuroimaging data processing framework in python. Frontiers in Neuroinformatics 5:13. 10.3389/fninf.2011.00013

Gorgolewski KJ, Esteban O, Markiewicz CJ, et al (2018) Nipype. Software 10.5281/zenodo.596855

Greve DN, Fischl B (2009) Accurate and robust brain image alignment using boundary-based registration. NeuroImage 48(1):63–72. 10.1016/j.neuroimage.2009.06.060

Hagberg AA, Schult DA, Swart PJ (2008) Exploring network structure, dynamics, and function using networkx. In: Varoquaux G, Vaught T, Millman J (eds) Proceedings of the 7th Python in Science Conference, Pasadena, CA USA, pp 11 – 15

Harte SE, Harris RE, Clauw DJ (2018) The neurobiology of central sensitization. Journal of Applied Biobehavioral Research 23(2):e12137

Hashmi JA, Baliki MN, Huang L, et al (2013) Shape shifting pain: chronification of back pain shifts brain representation from nociceptive to emotional circuits. Brain 136(9):2751–2768

Hotelling H (1992) Relations between two sets of variates. In: Breakthroughs in statistics: methodology and distribution. Springer, p 162–190

Hue TF, Lotz JC, Zheng P, et al (2024) Design of the comeback and backhome studies, longitudinal cohorts for comprehensive deep phenotyping of adults with chronic low-back pain (clbp): a part of the bacpac research program. medRxiv

Huntenburg JM (2014) Evaluating nonlinear coregistration of BOLD EPI and t1w images. Master’s thesis, Freie Universität, Berlin, URL http://hdl.handle.net/11858/00-001M-0000-002B-1CB5-A

Jenkinson M, Bannister P, Brady M, et al (2002) Improved optimization for the robust and accurate linear registration and motion correction of brain images. NeuroImage 17(2):825–841. 10.1006/nimg.2002.1132, URL https://www.sciencedirect.com/science/article/pii/S1053811902911328

Kamper SJ, Apeldoorn AT, Chiarotto A, et al (2014) Multidisciplinary biopsychosocial rehabilitation for chronic low back pain. Cochrane Database of Systematic Reviews (9)

Klein A, Ghosh SS, Bao FS, et al (2017) Mindboggling morphometry of human brains. PLOS Computational Biology 13(2):e1005350. 10.1371/journal.pcbi.1005350, URL http://journals.plos.org/ploscompbiol/article?id=10.1371/journal.pcbi.1005350

Konno Si, Sekiguchi M (2018) Association between brain and low back pain. Journal of Orthopaedic Science 23(1):3–7

Kregel J, Meeus M, Malfliet A, et al (2015) Structural and functional brain abnormalities in chronic low back pain: a systematic review. In: Seminars in arthritis and rheumatism, Elsevier, pp 229–237

Kwong J, Lin J, Leriche R, et al (2023) Quantifying pain location and intensity with multimodal pain body diagrams. JoVE (Journal of Visualized Experiments) (197):e65334

Lanczos C (1964) Evaluation of noisy data. Journal of the Society for Industrial and Applied Mathematics Series B Numerical Analysis 1(1):76–85. 10.1137/0701007, URL http://epubs.siam.org/doi/10.1137/0701007

Li X, Meng F, Huang W, et al (2024) The alterations in the brain corresponding to low back pain: Recent insights and advances. Neural Plasticity 2024(1):5599046

Maher C, Underwood M, Buchbinder R (2017) Non-specific low back pain. The Lancet 389(10070):736–747

Mansour A, Baria AT, Tetreault P, et al (2016) Global disruption of degree rank order: a hallmark of chronic pain. Scientific reports 6(1):34853

Mihalik A, Chapman J, Adams RA, et al (2022) Canonical correlation analysis and partial least squares for identifying brain–behavior associations: A tutorial and a comparative study. Biological Psychiatry: Cognitive Neuroscience and Neuroimaging 7(11):1055–1067

Power JD, Mitra A, Laumann TO, et al (2014) Methods to detect, characterize, and remove motion artifact in resting state fmri. NeuroImage 84(Supplement C):320–341. 10.1016/j.neuroimage.2013.08.048, URL http://www.sciencedirect.com/science/article/pii/S1053811913009117

Pruim RHR, Mennes M, van Rooij D, et al (2015) Ica-AROMA: A robust ICA-based strategy for removing motion artifacts from fmri data. NeuroImage 112(Supplement C):267–277. 10.1016/j.neuroimage.2015.02.064, URL http://www.sciencedirect.com/science/article/pii/S1053811915001822

Satterthwaite TD, Elliott MA, Gerraty RT, et al (2013) An improved framework for confound regression and filtering for control of motion artifact in the preprocessing of resting-state functional connectivity data. NeuroImage 64(1):240–256. 10.1016/j.neuroimage.2012.08.052, URL http://linkinghub.elsevier.com/retrieve/pii/S1053811912008609

Scherrer KH, Ziadni MS, Kong JT, et al (2021) Development and validation of the collaborative health outcomes information registry body map. Pain reports 6(1):e880

Schwenkreis P, Scherens A, Rönnau AK, et al (2010) Cortical disinhibition occurs in chronic neuropathic, but not in chronic nociceptive pain. BMC neuroscience 11:1–10

Seminowicz DA, Wideman TH, Naso L, et al (2011) Effective treatment of chronic low back pain in humans reverses abnormal brain anatomy and function. Journal of Neuroscience 31(20):7540–7550

Shen W, Tu Y, Gollub RL, et al (2019) Visual network alterations in brain functional connectivity in chronic low back pain: A resting state functional connectivity and machine learning study. NeuroImage: Clinical 22:101775

Shraim MA, Masśe-Alarie H, Farrell MJ, et al (2024) Neuroinflammatory activation in sensory and motor regions of the cortex is related to sensorimotor function in individuals with low back pain maintained by nociplastic mechanisms: A preliminary proof-of-concept study. European Journal of Pain 28(9):1607–1626

Treiber JM, White NS, Steed TC, et al (2016) Characterization and correction of geometric distortions in 814 diffusion weighted images. PLOS ONE 11(3):e0152472. 10.1371/journal.pone.0152472, URL http://journals.plos.org/plosone/article?id=10.1371/journal.pone.0152472

Tustison NJ, Avants BB, Cook PA, et al (2010) N4itk: Improved n3 bias correction. IEEE Transactions on Medical Imaging 29(6):1310–1320. 10.1109/TMI.2010.2046908

Wang HT, Smallwood J, Mourao-Miranda J, et al (2020) Finding the needle in a high-dimensional haystack: Canonical correlation analysis for neuroscientists. NeuroImage 216:116745

Wang S, Peterson DJ, Gatenby JC, et al (2017) Evaluation of field map and nonlinear registration methods for correction of susceptibility artifacts in diffusion MRI. Frontiers in Neuroinformatics 11. 10.3389/fninf.2017.00017, URL http://journal.frontiersin.org/article/10.3389/fninf.2017.00017/full

Wu A, March L, Zheng X, et al (2020) Global low back pain prevalence and years lived with disability from 1990 to 2017: estimates from the global burden of disease study 2017. Annals of translational medicine 8(6)

Zhang Y, Brady M, Smith S (2001) Segmentation of brain MR images through a hidden markov random field model and the expectation-maximization algorithm. IEEE Transactions on Medical Imaging 20(1):45–57. 10.1109/42.906424

Zhuang X, Yang Z, Cordes D (2020) A technical review of canonical correlation analysis for neuroscience applications. Human brain mapping 41(13):3807–3833

